# Color depth MIP mask search: a new tool to expedite Split-GAL4 creation

**DOI:** 10.1101/318006

**Authors:** Hideo Otsuna, Masayoshi Ito, Takashi Kawase

## Abstract

The GAL4-UAS system has proven its versatility in studying the function and expression patterns of neurons the *Drosophila* central nervous system. Although the GAL4 system has been used for 25 years, recent genetic intersectional tools have enabled genetic targeting of very small numbers of neurons aiding in the understanding of their function. This split-GAL4 system is extremely powerful for studying neuronal morphology and the neural basis of animal behavior. However, choosing lines to intersect that have overlapping patterns restricted to one to a few neurons has been cumbersome. This challenge is now growing as the collections of GAL4 driver lines has increased. Here we present a new method and software plug-in for Fiji to dramatically improve the speed of querying large databases of potential lines to intersect and aid in the split-GAL4 creation. We also provide pre-computed datasets for the Janelia GAL4 (5,738 lines) and VT GAL4 (7,429 lines) of the Drosophila central nervous system (CNS). The tool reduced our split-GAL4 creation effort dramatically.

## Introduction

Genetic targeting and control of defined sets of neurons in transgenic animals has accelerated our understanding of the morphology and function of the nervous system in many animals. This has been especially apparent in the fruit fly, *Drosophila melanogaster*. In flies, the GAL4-UAS system is a robust method for targeting gene expression into specific neurons (Brand and Perrimon, 1993; Jenett et al., 2012; Pfeiffer et al., 2008). Many large collections of GAL4 driver lines are publically available (Dionne et al., 2018; Jenett et al., 2012; Pfeiffer et al., 2008; Tirian and Dickson, 2017; http://flweb.janelia.org/cgi-bin/flew.cgi), and although these collections have yielded many new insights into the neural basis of behavior (Otsuna et al., 2014; Robie et al., 2017) the pattern of neuronal expression remains broad in these lines making it difficult to ascribe function to individual cells. Further refinement of the expression patterns is possible using the split-GAL4 system is used to eliminate unnecessary GAL4 expression from the out of target neurons and achieve individual cell-type specificity (Luan et al., 2006).

To achieve this specificity, activation domain (AD) and DNA binding domain (DBD) hemidriver ‘split-halves’ that are derived from GAL4 patterns are combined and drive the expression of the GAL4 system only in overlapping cells (for a detailed review see Dionne et al., 2018). This intersectional approach to generating split-GAL4 driver lines is a powerful tool in Drosophila neurobiology (Aso et al., 2014; Namiki et al., 2017; Wu et al., 2016), but the screening of confocal stacks in 3D to find overlapping neurons from more than ten thousand of GAL4 lines is an onerous task on the part of the researcher. Here we present a novel way to rapidly query the expression patterns of thousands of driver lines and computationally determine which lines likely contain a neuron or set of neurons of interest. We do this by first generating maximum intensity projection images (MIPs) where color represents the depth.

Thus, the neuron searching method for the split-GAL4 creation is ideal due to the following characteristics: it is non-segmentation based, the original x-y resolution is maintained, it not only queries for neurons but also can provide other neuronal expression levels, the original brightness of the neurons is reflected, there is an easily modified 3D mask, and the ease of adding additional datasets into the searching data. Therefore, this searching method is not just for individual neurons, but can query entire expression patterns as seen in GAL4 driver lines.

Here are our results for the new 3D neuron searching method for the split-GAL4 creation from original GAL4 expressed data.

## Results

Our searching method creates a matching score from the overlapped pixel number of color depth MIP (Timo et al,, 2006). The pixel color represents z-depth. The xy resolution is same as the JFRC2010 template. If the color and xy position of the pixel match between the mask and the searching data, then the approach here will count it as a positive matching score. The matching color range can be set to certain fluctuation; this will adjust for alignment errors and individual brain/neuronal shape difference between samples. Unlike the skeleton, our non-segmentation based searching method can use the original thickness of the neurons. Typically, a single neuron has around 5 pixel thickness in our JFRC2010 aligned brain. This thickness also adjusts for neuronal position difference between samples.

## Workflow

Original confocal stacks need to be aligned to a single 3D template. Then the color depth MIPs are generated from each of aligned stacks individually (Fig. 1A-D). If there are the same neurons in different samples; the MIP color of the neurons became almost identical after the alignment step. To search for a neuron of interest, the user can create a mask by using the freehand tool in Fiji (Fig. 1E). The search is performed against the color MIP dataset; the positive hit is usually 0.5~2% of the original dataset (Fig.1 F).

**Figure 1.**
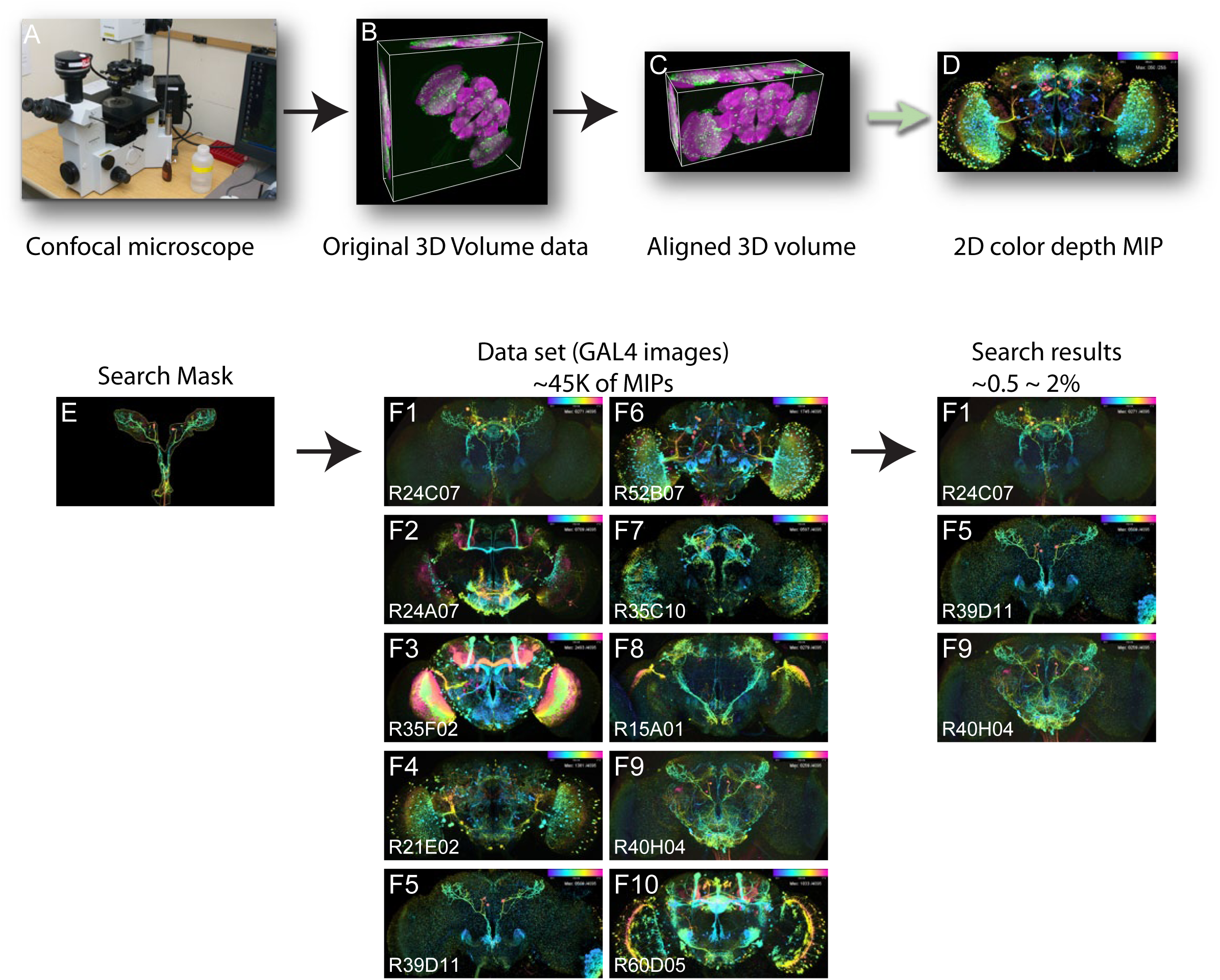
Workflow. **A-C:** Original confocal stacks need to be aligned to a single 3D template. **D:** The color depth MIP is generated from the aligned stacks. **E:** Example of a searching neuron. **F:** example of searching dataset and the results. The search results contain the target neuron as a positive hit.

## Automatic-brightness adjustment

Before the MIP creation, our program adjusts the image brightness automatically for enhancing the neuron searching sensitivity. To set neuronal brightness equally between different brains, we create the mask for the neuron fibers only but not for the cell bodies/tick neuron bundles (Fig. 2A, B). We employed the 2D rod shape neuronal segmentation fitting method to the neuronal fiber segmentation - Direction selective local thresholding (DSLT) (Kawase et al., 2015). The method allows us to avoid the intensity measuring of the bright bold structures. The adjusted brightness is for fine neuronal fibers. Our program adjusts averaged pixel value to desired brightness value (usually 190~200 /255 gray value) within the 2D mask. The dimmer signal becomes brighter after using high-thresholding of the gray value. Though that approach, the user can tell the brightness of the sample from this thresholding value. The dimmer samples require more brightness enhancement with lower thresholding value. But the images from different brains will look the same for brightness (Fig 2C, C’, D).

**Figure 2.**
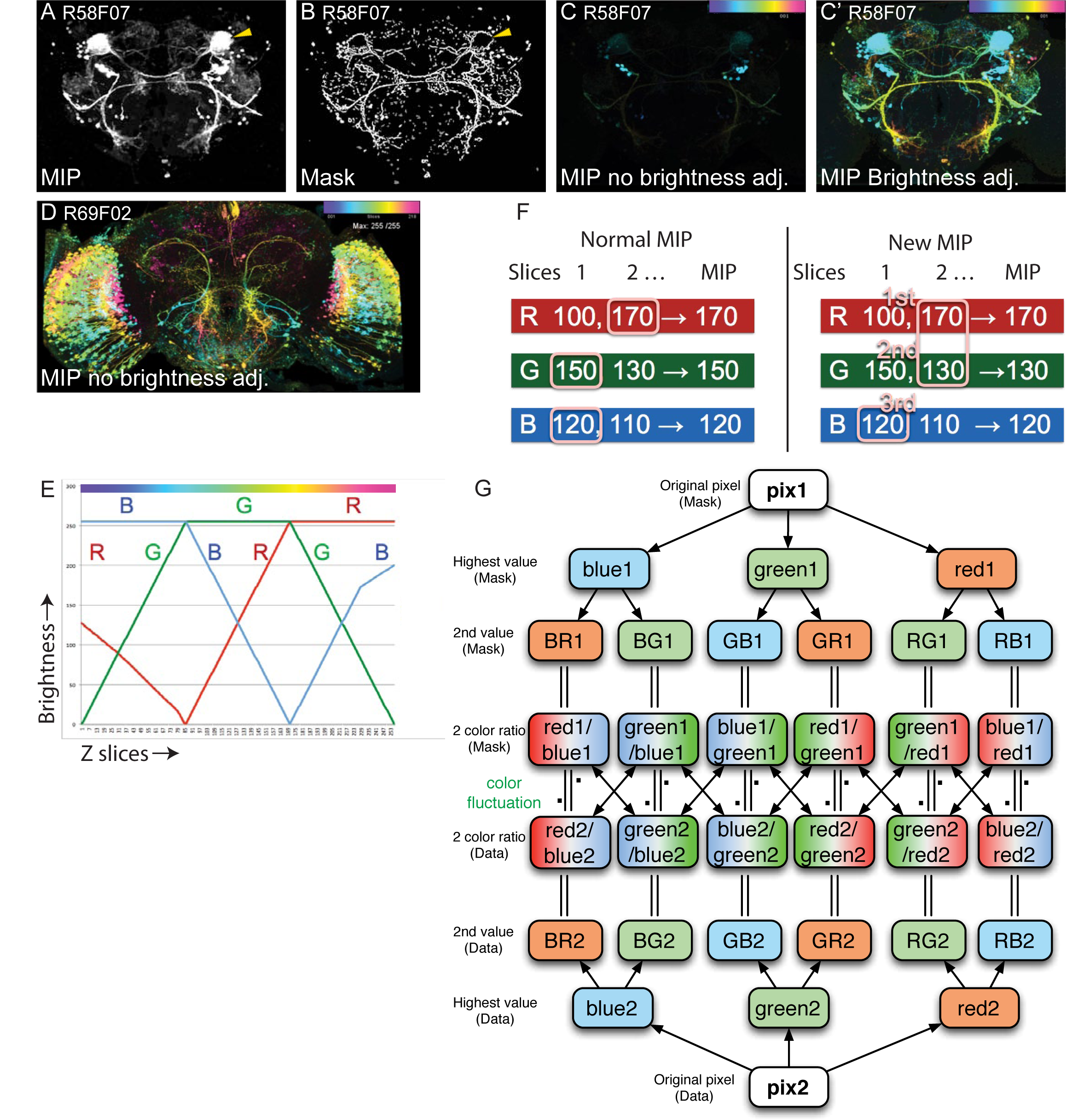
Color depth MIP creation and the mask search algorithm. **A-B:** The mask creation for the brightness measurement of neuron fibers. **A:** Original MIP signal, **B:** Direction sensitive local threshold (DSLT) mask. The mask skips bold and bright structures (Arrow: the elipsoid body). Thus, the measured brightness is mainly from the neuron fibers. **C-D:** Example of the auto-brightness adjustment. **C:** Original image of the R58F07 line. The GAL4 expression is too weak and could not show the neuro-fibers. **C’:** Auto-re-mapped the gray scale value 20 as 255 from the C. **D:** The strong GAL4 expression line R69F02, the brightness of the image is original value. C’ and D are almost similar brightness. **E:** Color look up table of our color depth MIP. Y axis is the brightness of the signal. X axis is Z-slice number. One of the RGB channels represents original gray scale value (Top line, 1st value). The ratio between 1st and 2nd color value represents unique Z-slice number within the 255. **F:** The MIP method comparison between normal MIP and our MIP. In the normal MIP (left panel), the RGB signals between different slices are independent. Just the maximum value of each RGB will create the MIP. The MIP will create the 1^st^ value (R-170) and the 2^nd^ value (G-150) pair. The pair does not exist within the z-stack. In our MIP (right panel), the 1st value and the 2nd value are paired from a single slice. **G:** Color matching algorithm between the mask and the searching data.

## Z-depth Color MIP creation

For the accurate single slice z-resolution and the easy visual screening of the results, we created new color look-up-table (LUT) (Fig. 2E). We also modified the method of MIP creation to keep the accurate z-depth information in the MIP (Fig. 2F). The color matching algorithm is based on our LUT. Our method counts only the top two gray values among three of RGB channels for the color matching. The highest gray value represents original gray value. The ratio between 1^st^ and 2^nd^ top gray values represents the z-slice number. If the 3^rd^ gray value is lower then the 2^nd^ gray value, then the highest 3^rd^ value will be accepted as a MIP signal. We found that the 3^rd^ value enhanced the visual screening efficiency (Fig. 2G).

## Mask creation

The screening process requires quick 3D mask creation and quick result image browsing. We implemented the program in Fiji (https://fiji.sc/). The user can create mask from either 3D segmented neuron or the color depth MIP image. If the former method is applied, we have been developing branch out version of the FluoRender software (Wan et al., 2009), VVDviewer (https://github.com/takashi310/VVD_Viewer). The software provides the user with quick 3D segmentation on the 2D display. If the latter method is applied, the user can use the Fiji’s 2D ROI creation function (Polygon/Freehand selections). Either way, the 3D mask creation will only need a few minutes.

## Searching algorithm and speed

Color matching algorithm between the mask and the searching data are in Fig. 2G. The pixel from mask: pix1, the pixel from the data: pix2. Both pixels create “two color ratio” from the highest color value (1^st^ value) and 2^nd^ color value. Center of the images: The color depth MIP mask search plugin compares between these two color ratios. Higher color fluctuation setting will accept wider differences between the ratios: it allows the color matching between more z-depth shifting, as well as x-y displacements of up to 4 pixels to compensate for differences in alignment between samples.

The overlap search is performed only within the mask area. Thus, the searching is efficient and fast. If the user loads images from SSD performance is ~15 seconds for 8,000 samples 11.7GB, ~30-45 seconds from the HDD, ~7 seconds from the system memory.

## Reviewing search results

We also found that the reviewing process of searching result is critical in the workflow. If the user needs to spend time more than few seconds per image, it is not intuitive. For example, the original searching images are ~46,445, then the mask search will narrow down to ~1% (200~500) of images. Usually, the user needs to check top 50~150 images. Our color depth MIP visualization allows the user to check the 50~150 images within ~10 minute. Color depth MIP mask search method itself could not hit the line if the targeted neuron is covered by the cloud of neurons. This feature matches to both of the AD/DBD split-GAL4 picking and the reviewing process.

## Comparison with NBLAST

For adding new aligned data to an exiting database, NBLAST (Costa et al., 2016) needs 3D segmentation and conversion to the vector data by using R coding. It then needs to associate the NBLAST data and GAL4 line MIPs in order to make it effective for the split-GAL4 to be picked-up. However, in our color depth MIP method, users only need to create the MIP and move the file into an existing MIP set folder. Unlike other searching databases, the color depth MIP searching method is searching the original GAL4 image itself. It does not require linking to the image and searching query.

## Search results

We compared the search results by using ~3500 of Janelia GAL4 lines (http://flweb.janelia.org/cgi-bin/flew.cgi). We used three dataset for our comparison: this Color depth MIP, the NBLAST dataset from VirtualFlyBrain (VFB) and the NBLAST dataset with newly DSLT segmentation. The NBLAST data from the VFB missed many of neurons (Fig. 3B). Thus, we re-aligned the Janelia Gen1 GAL4 lines and newly 3D segmented GAL4 expressed neurons by the DSLT high-sensitive 3D segmentation method (Fig. 3C). Then we created the new NBLAST database. This new dataset contains many more segmented neurons than the VirtualFlyBrain’s dataset. The test searching masks are generated from previously published split-GAL4 lines (http://splitgal4.janelia.org/cgi-bin/splitgal4.cgi, R_MB027B, R_OL0001B (LC22), R_OL0015B (LC11), SS0732, SS4159 (Aso et al., 2014; Shigehiro Namiki, 2017; Wu et al., 2016). We choose three different neuronal shape characters for the searching test. Bold bright fiber: R_MB027B, tick fascicle lobula columnar neuron: R_OL0001B (LC22), R_OL0015B (LC11) (Otsuna and Ito, 2006), fine neuro-fiber: SS0732, SS4159.

**Figure 3.**
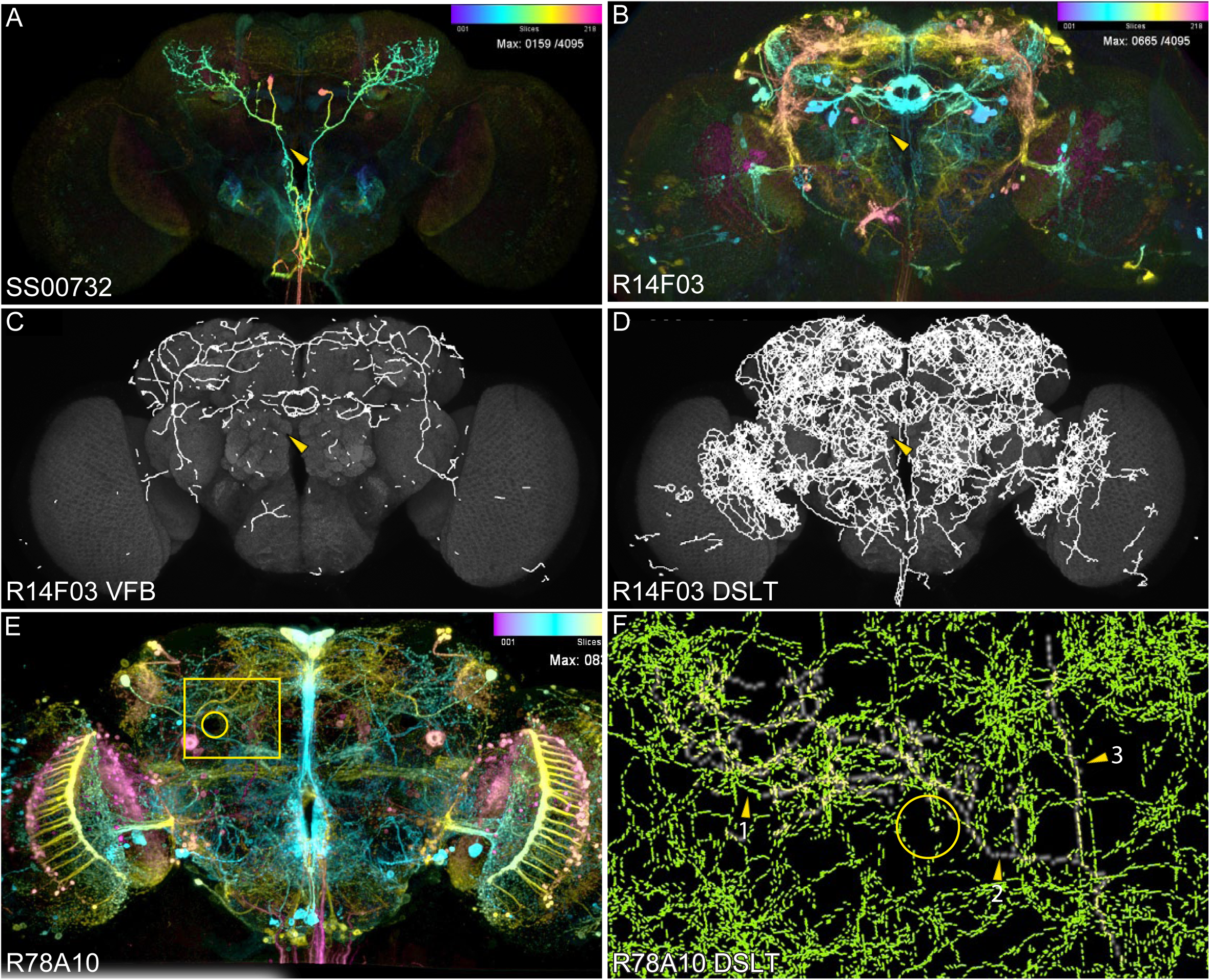
The analysis of neuron search results. **A:** Aimed neuron SS00732 split-GAL4 line, **B:** The color MIP of R14F03 GAL4 line. The line was used as AD line. The color MIP mask search hits this line as rank #23 and the R24C07 (DBD) line as rank #4. **C:** NBLAST VFB could not hit the R14F03 line due to lack of the segmentation (Yellow arrow head). But it hits R24C07 as rank #3. **D:** NBLAST DSLT hits the R14F03 line as rank #17 but could not hit the R24C07 line within the top 50. **E-F:** Top score GAL4 line R78A10 in the NBLAST DSLT. The line has too much GAL4 expression. It appears to be false hit. In fact, it is due to increase score by high density of other fibers. It is difficult to tell if this is real hit or not. **E:** Original GAL4 expression. The yellow rectangle area in F. Yellow circle is the position of mushroom body peduncles. **F:** Green: local vector, White: Submitted skeleton. Arrow head (1): The area has dense arborization from other neurons. some local vectors are matching with SS00732 neuron. (2): local vectors are not existing in the trunk, (3): the vertical line is the cell body fiber. Many other neurons have cell body fiber in the area.

In our test, NBLAST tends to hit high-density GAL4 expressed line (Table 7 K-N, Table 8 H-N). These high-density GAL4 expressed lines are not desirable for the split-GAL4 creation. The NBLAST matching score is a result of the local point vector matching. Surrounding neurons could increase matching score (Fig. 3E, F). Thus, our new high-sensitive segmented NBLAST dataset ranked the specific GAL4 expressed lines lower than the high-density GAL4 expressed lines (Tables 1, 3, 4, 5).

**Table 1.**
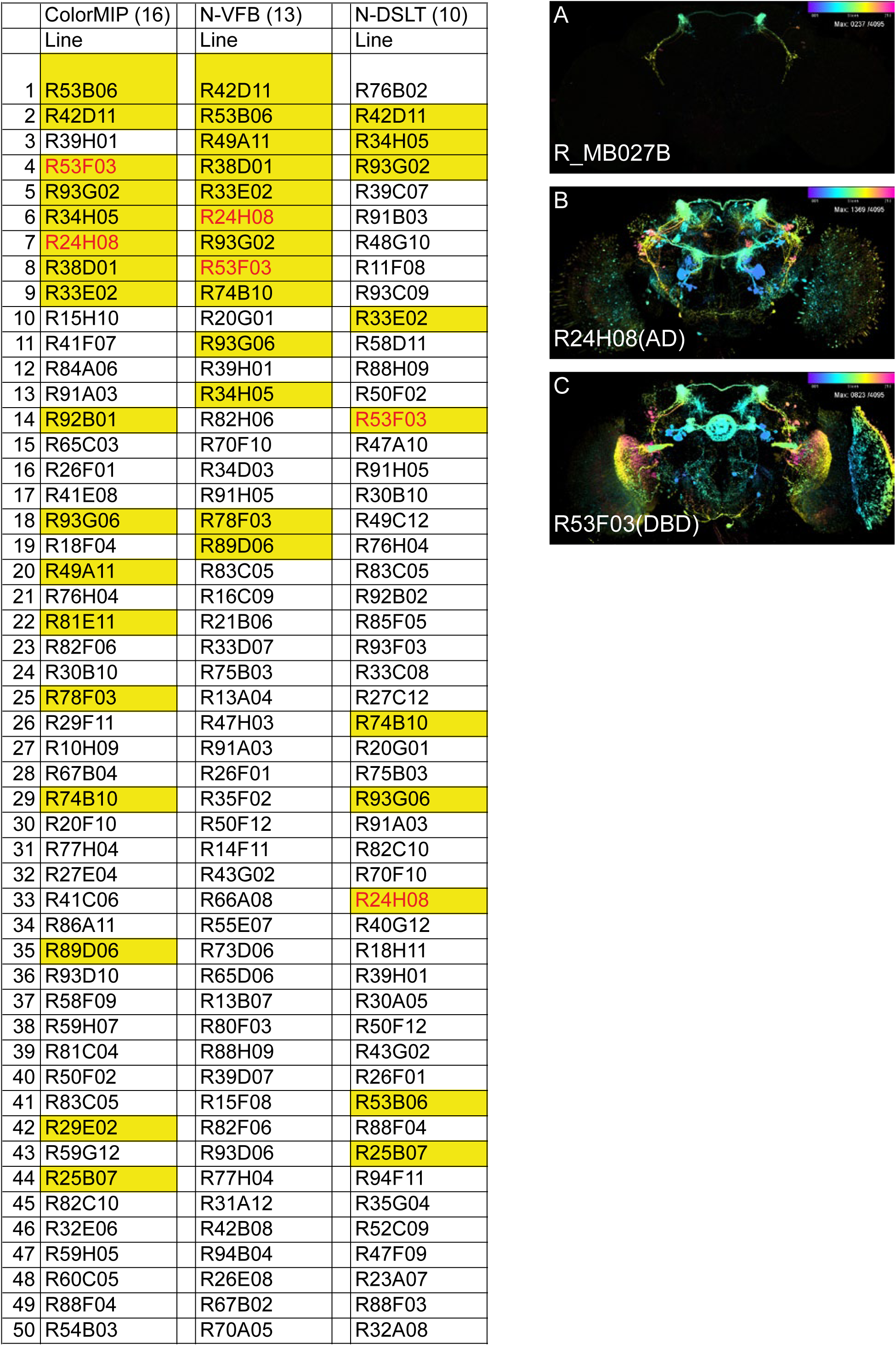
Searching results of a neuron from the stable split-GAL4 line. The tables are the top 50 hits from multiple search methods: from the left, Color depth MIP search, NBLAST data from virtual fly brain, NBLAST data from our high-sensitive DSLT segmentation. All searching methods have the same Janelia GAL4 dataset. In the table, yellow cell represents real hit of the neuron. The yellow cell + red character represents original AD/DBD line. A: The searched neuron from the split-GAL4 line. B: The GAL4 line image used as AD. C: The GAL4 line image used as DBD.

**Table 2.**
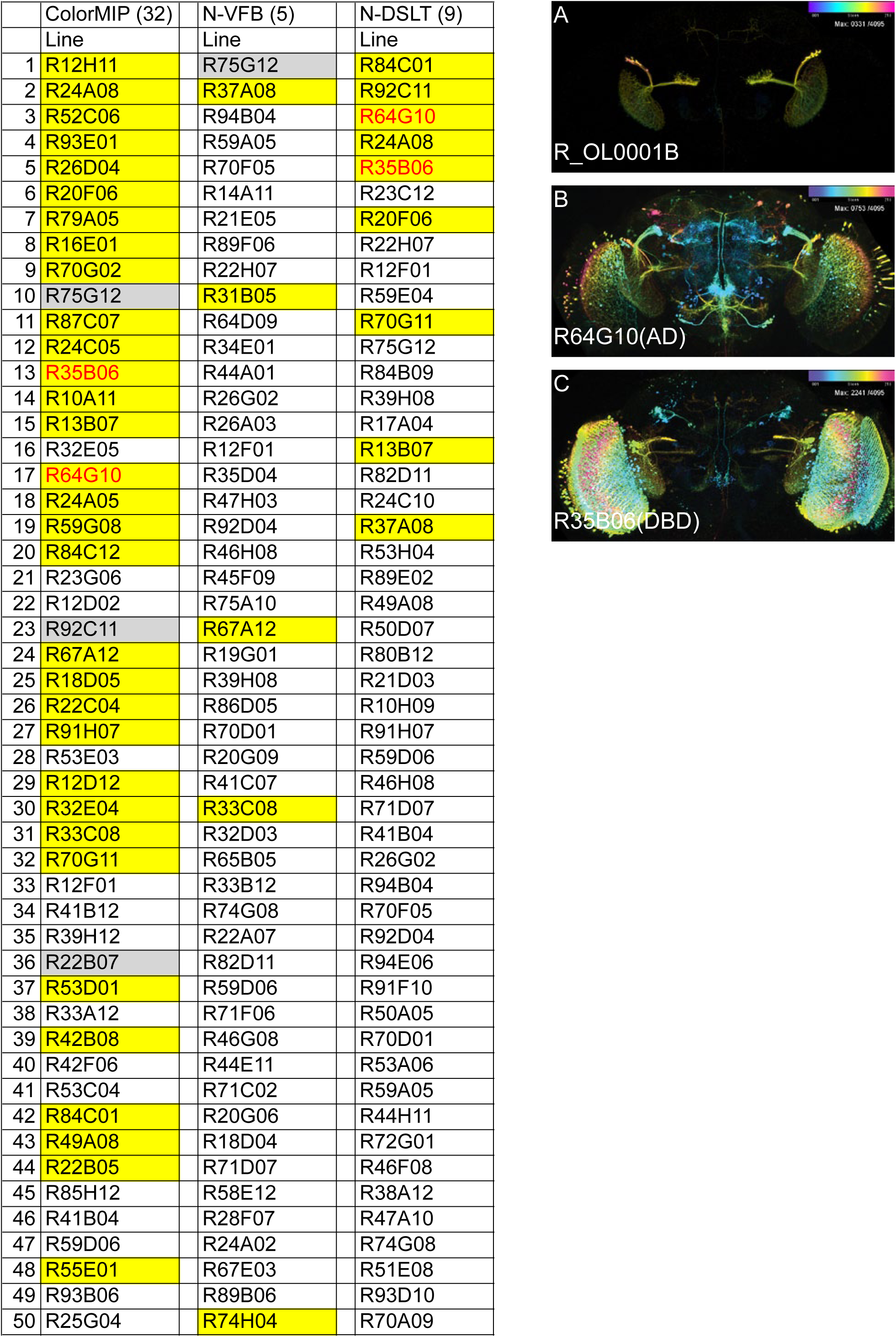
Searching results of a neuron from the stable split-GAL4 line. The tables are the top 50 hits from multiple search methods: from the left, Color depth MIP search, NBLAST data from virtual fly brain, NBLAST data from our high-sensitive DSLT segmentation. All searching methods have the same Janelia GAL4 dataset. In the table, yellow cell represents real hit of the neuron. The yellow cell + red character represents original AD/DBD line. A: The searched neuron from the split-GAL4 line. B: The GAL4 line image used as AD. C: The GAL4 line image used as DBD.

**Table 3.**
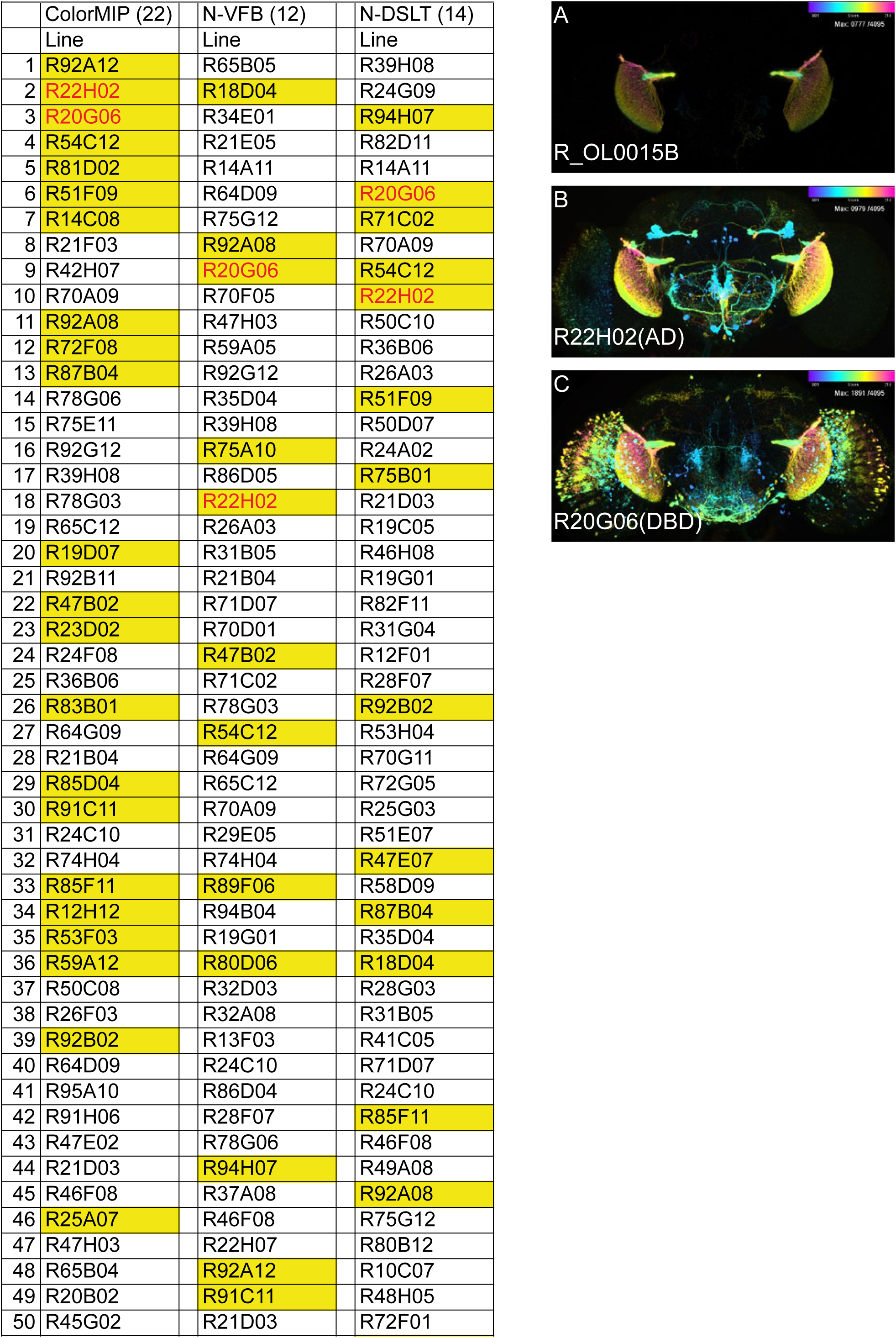
Searching results of a neuron from the stable split-GAL4 line. The tables are the top 50 hits from multiple search methods: from the left, Color depth MIP search, NBLAST data from virtual fly brain, NBLAST data from our high-sensitive DSLT segmentation. All searching methods have the same Janelia GAL4 dataset. In the table, yellow cell represents real hit of the neuron. The yellow cell + red character represents original AD/DBD line. A: The searched neuron from the split-GAL4 line. B: The GAL4 line image used as AD. C: The GAL4 line image used as DBD.

**Table 4.**
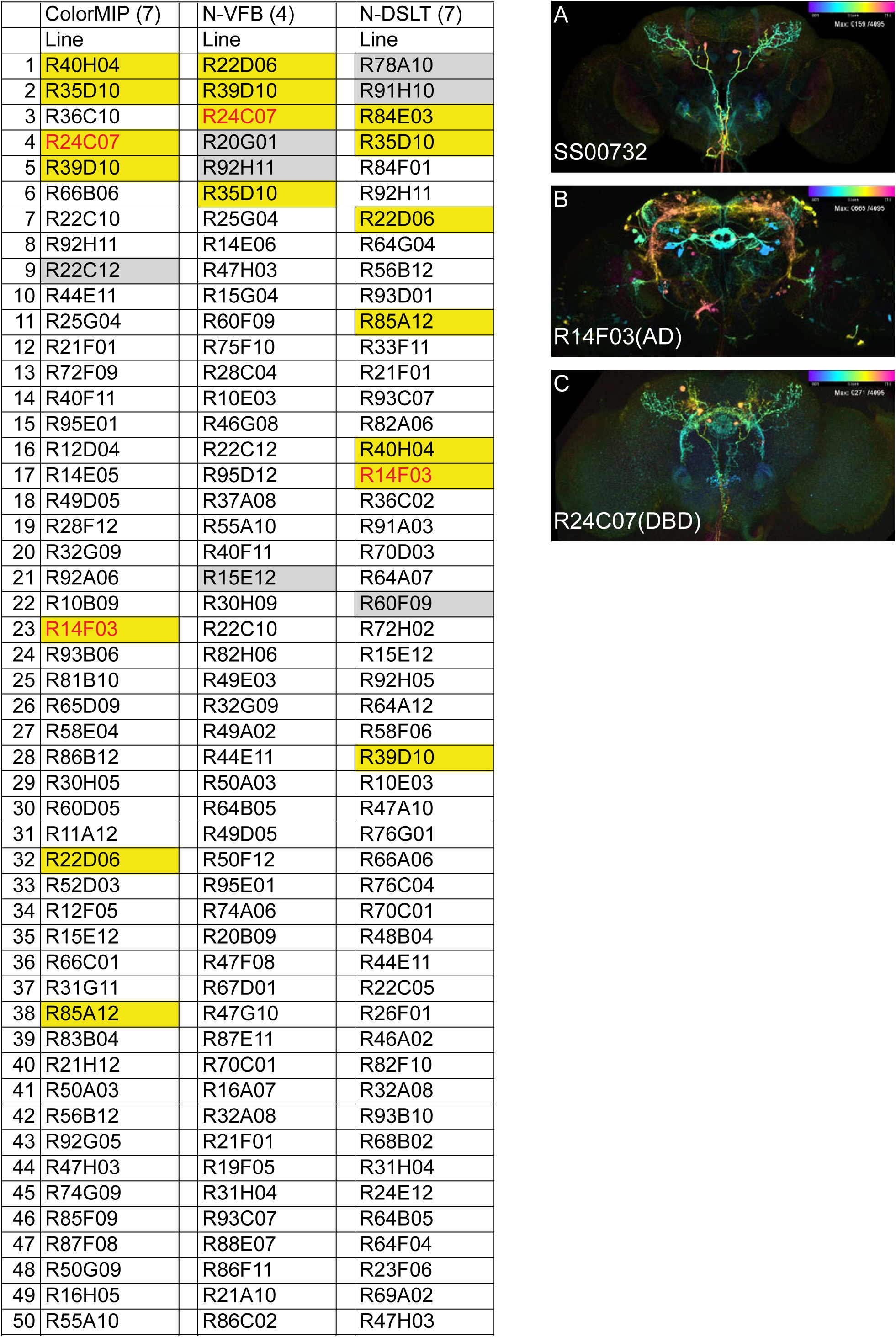
Searching results of a neuron from the stable split-GAL4 line. The tables are the top 50 hits from multiple search methods: from the left, Color depth MIP search, NBLAST data from virtual fly brain, NBLAST data from our high-sensitive DSLT segmentation. All searching methods have the same Janelia GAL4 dataset. In the table, yellow cell represents real hit of the neuron. The yellow cell + red character represents original AD/DBD line. A: The searched neuron from the split-GAL4 line. B: The GAL4 line image used as AD. C: The GAL4 line image used as DBD.

**Table 5.**
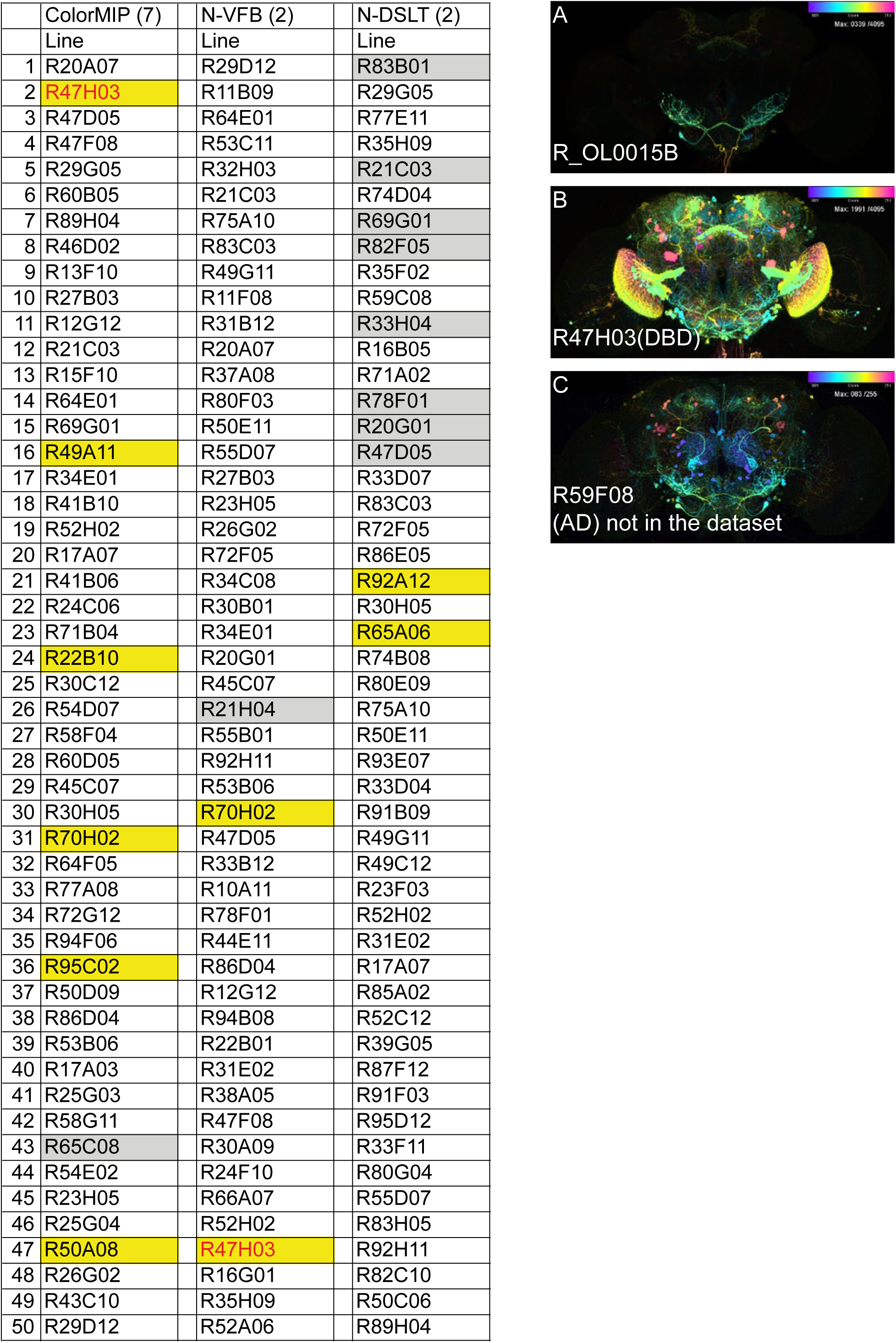
Searching results of a neuron from the stable split-GAL4 line. The tables are the top 50 hits from multiple search methods: from the left, Color depth MIP search, NBLAST data from virtual fly brain, NBLAST data from our high-sensitive DSLT segmentation. All searching methods have the same Janelia GAL4 dataset. In the table, yellow cell represents real hit of the neuron. The yellow cell + red character represents original AD/DBD line. A: The searched neuron from the split-GAL4 line. B: The GAL4 line image used as AD. C: The GAL4 line image used as DBD.

**Table 6.**
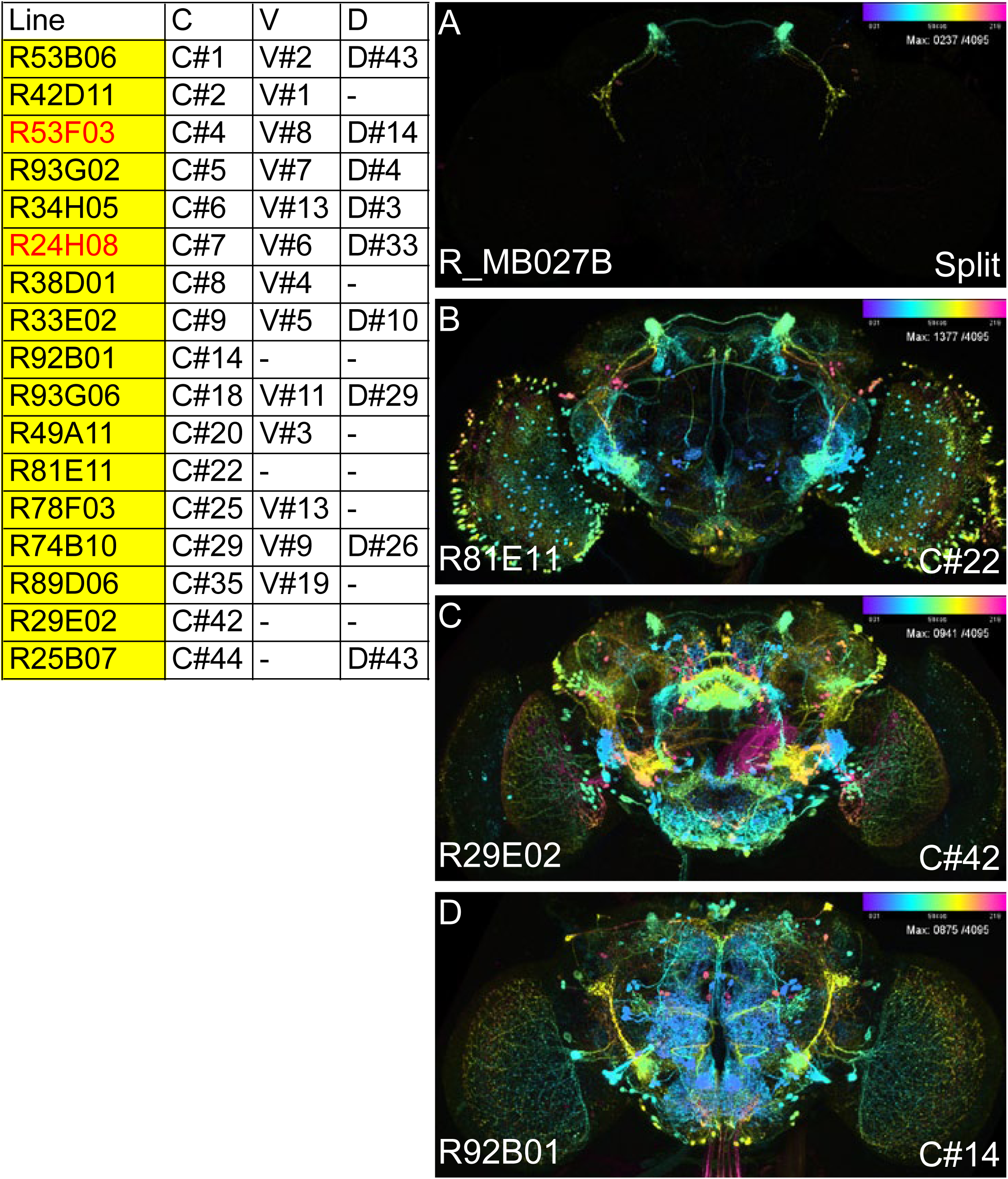
The comparison among the positive hits from three-searching methods. A: The neuron used as searching mask from the original split-GAL4 line. B-N: The positive hits GAL4 images that contain the neuron from split-GAL4 lines. C#: hits by color depth MIP search, the number is ranking. V#: hits by NBLAST search by using virtual fly brain data. D#: NBLAST search hits by using DSLT segmented data.

**Table 7.**
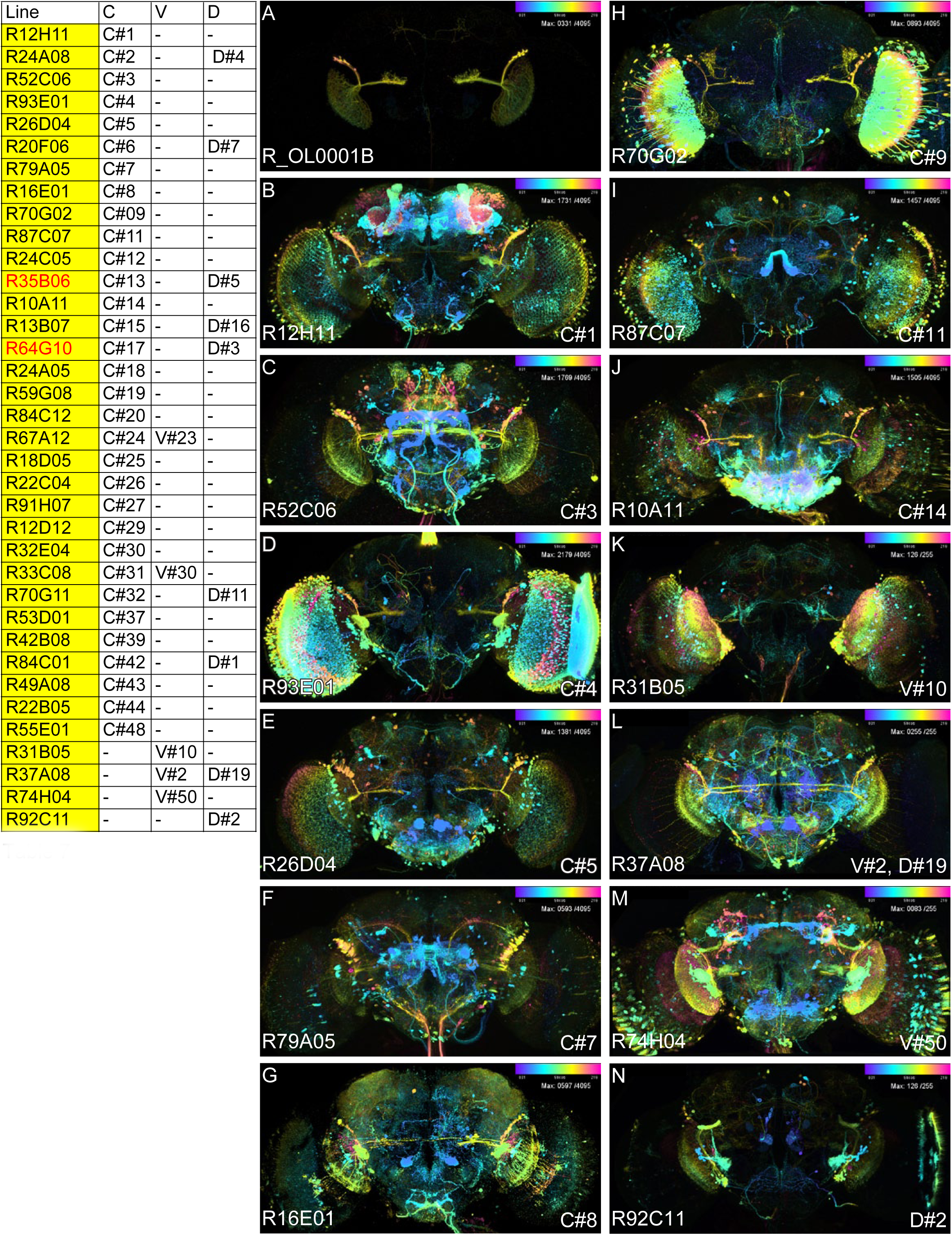
The comparison among the positive hits from three-searching methods. A: The neuron used as searching mask from the original split-GAL4 line. B-N: The positive hits GAL4 images that contain the neuron from split-GAL4 lines. C#: hits by color depth MIP search, the number is ranking. V#: hits by NBLAST search by using virtual fly brain data. D#: NBLAST search hits by using DSLT segmented data.

**Table 8.**
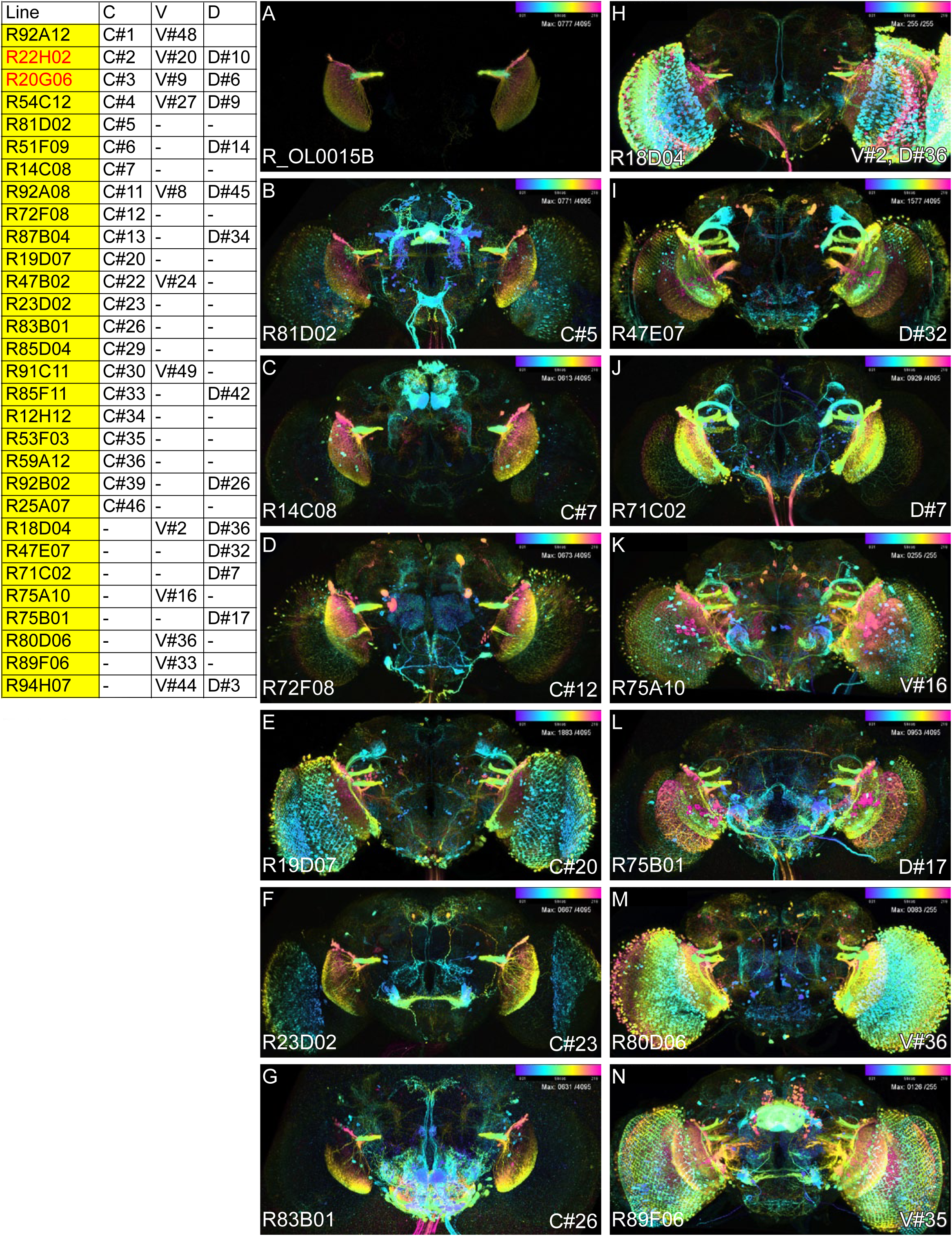
The comparison among the positive hits from three-searching methods. A: The neuron used as searching mask from the original split-GAL4 line. B-N: The positive hits GAL4 images that contain the neuron from split-GAL4 lines. C#: hits by color depth MIP search, the number is ranking. V#: hits by NBLAST search by using virtual fly brain data. D#: NBLAST search hits by using DSLT segmented data.

NBLAST dataset from the VFB worked well only with the mushroom body output neurons from the R_MB027B split line within the five neurons test. The neuron was bright and bold. It was segmented nicely in many cases. However, both of the lobula columnar neurons were not detected well by the NBLAST VFB (Table 2, 3), because the segmentation of the lobula are often lacking. This is the reason of the NBLAST DSLT containing more hits than the NBLAST VFB of the lobula columnar neurons (Table 2, 3). However, The NBLAST DSLT was too sensitive to any of the lobula columnar neurons. Therefore, the real hits are spread within top 50 ranking (Table 3) and less number of real hit (Table 2). The lack of segmentation of the NBLAST VFB is also observed in Table 4 and 5. In Table 4, NBLAST VFB could not hit R_14F03 (AD) line within top 50, due to the lack of segmentation (Fig. 3A-C).

The color depth MIP searching method could hit more useful samples for the split-GAL4 creation than the NBLAST VFB and NBLAST DSLT with both the fine fibers and bold LC neurons in our test (Table 6 - 10). For example, in Table 8, B-G are hit only by the color depth MIP searching, H-N are only hit by the NBLAST searching at more than top 50 ranks. The relatively specific LC11 GAL4 expression is in B, C, D, F, G but not in H, I, K, L, M, N.

**Table 9.**
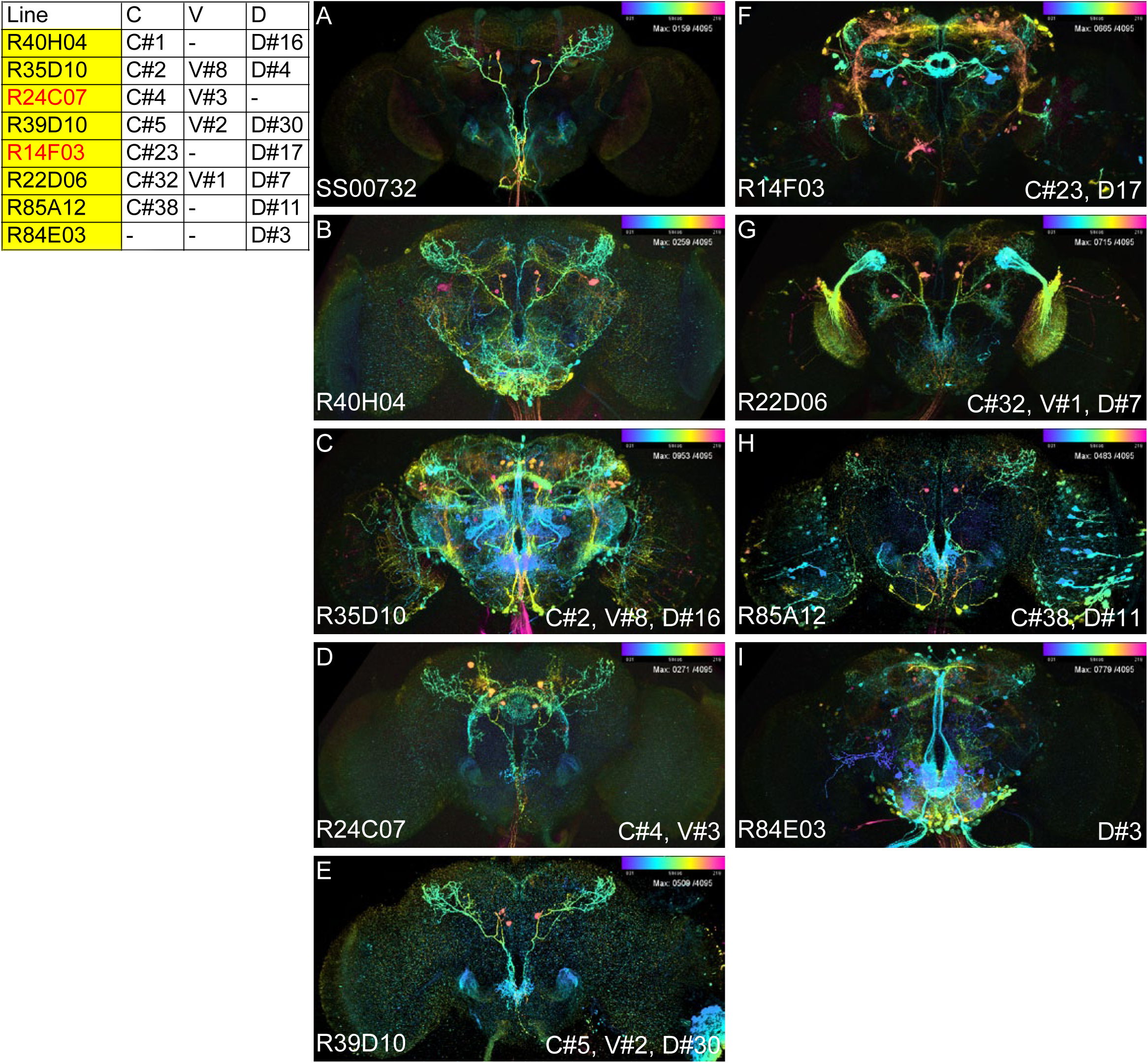
The comparison among the positive hits from three-searching methods. A: The neuron used as searching mask from the original split-GAL4 line. B-N: The positive hits GAL4 images that contain the neuron from split-GAL4 lines. C#: hits by color depth MIP search, the number is ranking. V#: hits by NBLAST search by using virtual fly brain data. D#: NBLAST search hits by using DSLT segmented data.

**Table 10.**
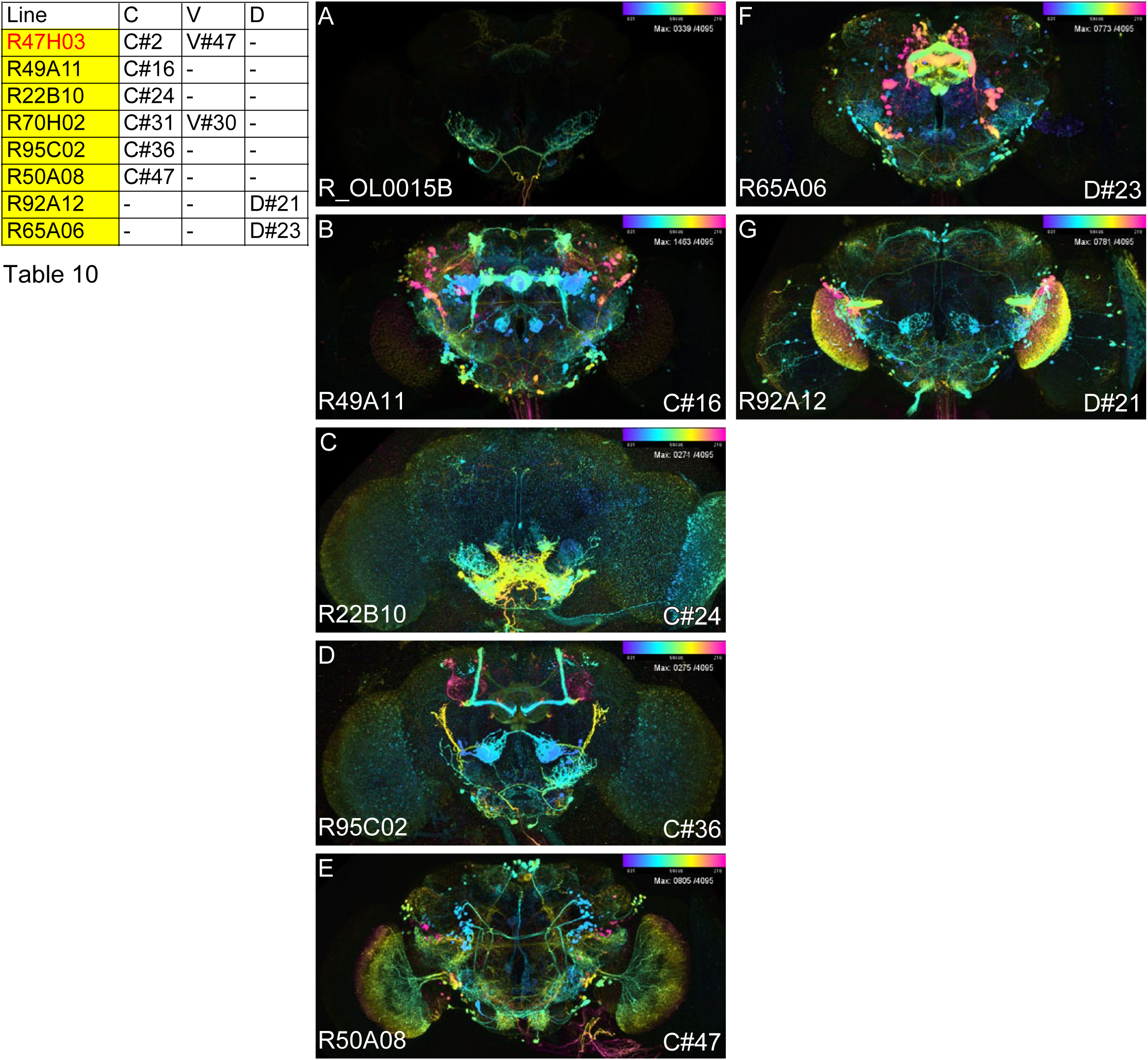
The comparison among the positive hits from three-searching methods. A: The neuron used as searching mask from the original split-GAL4 line. B-N: The positive hits GAL4 images that contain the neuron from split-GAL4 lines. C#: hits by color depth MIP search, the number is ranking. V#: hits by NBLAST search by using virtual fly brain data. D#: NBLAST search hits by using DSLT segmented data.

The high-sensitivity of the color depth MIP searching is based on the automatic brightness adjustment. For example, for line SS00732 the AD line (14F03, 665/4095 brightness increased, Table 6F) and the DBD line (24C07, 271/4095 brightness, Table 6D) are both so dim for the original GAL4 expression. Their brightness is increased significantly during the MIP creation.

## Resources from this study

The 46,445 color depth GAL4 brain MIPs from the Gen1-R GAL4 (5,738 lines) and VT GAL4 (7429 lines). The 9,395 of LexA brain MIPs from the Janelia Gen1-LexA (1,100 lines) and VT LexA (2020 lines). The 15,057 of GAL4 VNC MIPs from the Janelia Gen1-GAL4 (7,491 lines) and VT GAL4 (2795 lines). The 3,534 of LexA VNC MIPs from the Janelia Gen1-LexA (1,536 lines) and VT LexA (1,497 lines). The pre-computed color depth MIPs links to the source code can be found at: https://www.janelia.org/open-science/color-depth-mip

The Fiji based color depth MIP searching program is in this location (https://github.com/JaneliaSciComp/ColorMIP_Mask_Search). VVDviewer: 3D/4D visualization and 3D segmentation software (https://github.com/takashi310/VVD_Viewer).

## Discussion

From the searching test of the color depth MIP method, we could hit all of AD/DBD split pairs for all five split-GAL4 lines. (Split-GAL4 line: SS00732, R24C07 (DBD) is not in the NBLAST VFB dataset. However, when we added the color depth MIP to the searching dataset, it was found as the 40^th^ highest hit (Data not shown). This result suggests that our searching method is useful for split-GAL4 creation. Also, we have internally validated using the the color depth MIP method in multiple visitor projects and labs in Janelia (https://www.janelia.org/you-janelia/visiting-scientists/active-projects: 1) Generation of cell-type-specific GAL4 driver lines for neurons of the Drosophila adult leg neuropil; 2) Whole Fly Brain Tracing Effort: The Lobula Plate Tangential cells: elucidating the feed-forward and feedback circuitry; 3) Generation of cell-type specific GAL4 driver lines for neurons in the Drosophila adult flight and haltere neuropil of the ventral nerve cord; 4) Mapping and analyzing neurons in the terra incognita regions of the Drosophila brain; 5) Mapping the lateral horn of Drosophila; 6) Generation of cell-type specific GAL4 driver lines for neurons in the Drosophila adult suboesophageal ganglion; 7) Development of Hemilineage Tools for Studying Neurons of the Adult Drosophila CNS; 8) Genetic dissection of somatosensory neural circuits in the Drosophila ventral nerve cord). Our method is extremely useful for picking AD/DBD pairs from more than 46,445 Gen1 GAL4 images. Multiple researches can pick up ~90% of AD/DBD pairs that likely target the neuron(s) of interest. Then they could create split-GAL4 lines around 60~80% of the whole known neuron type (results will be published in future), though the individual success rate for a given intersection typically is in the 10-30% range as detailed in Dionne et al., 2018.

In order to accelerate neuron seaching in large datasets, the NBLAST and Braingazer programs were introduced (Bruckner et al., 2009; Costa et al., 2016). Though these tools are very valuable for specific applications, in our hands these methods weren’t well suited for split-GAL4 line creation. The split-GAL4 system requires identifying specific expression patterns in overlapping parental driver lines that have little to no overlap of off target neurons. Neither of these methods is accounts for other expression within the GAL4 line potentially leading to non-specific split-GAL4 expression. In addition, both methods require 3D neuronal segmentation to enable searching. Segmentation of high-density GAL4 expression patterns is challenging. Moreover, split-GAL4 driver lines are often better produced from strongly expressed parental lines. Previous methods do not account for the brightness, and thus expression level, of the neurons. Current NBLAST databases rely on skeletonization don’t account for the thickness of fascicles. If the mask is a single neuron, and target is a thick neuronal fascicle with a different shape, NBLAST has problems with detection. Since searching the GAL4 dataset is not a single neuron, the user often needs to modify the 3D mask to pick the targeted neuron. The 3D mask modification is also another challenge in NBLAST but not in Braingazer. However, Braingazer’s database provides averaged signals from three different aligned brains, which means fine fibers are easily missed from the database.

Through the analysis of mask search results, we found that many neurons have relatively high shape fluctuations in the neuronal trunk and location of the cell body, but not in the projections. Some neurons exhibit remarkable stability, especially those that aggregate in tracts. These stable characteristics are useful to produce high hit rates. Some neurons are easy to target, but some are not. So, if the neuron has high stable character, it is relatively easy to hit.

We analyzed the alignment fluctuations in the Janelia Gen1 GAL4 collection and found that neurons are usually within ~5 px in the JFRC2010 template resolution (1024 x 512 px). Since most neurons are 3~5 px thick, the alignment fluctuation is within the range of neuron fiber thickness. But if we use skeletonized neurons for searching, the alignment variability will be a challenge.

In order to eliminate high density GAL4 expressed lines, our color depth MIP method allows the user to skip pre-calculation of the density of expression from many brain areas. The MIP itself shows the strongest GAL4 expressed neurons as a result. MIP generation will not render the targeted neuron if the expression pattern around the targeted neuron is too dense, thus excluding patterns that are too dense. Therefore the result of the color depth MIP search contains more specific GAL4 lines, meaning the color depth MIP searching method is very useful for split-GAL4 AD/DBD selection.

## Method

We used aligned confocal images of the Janelia and VT GAL4 collections (Jenett et al., 2012; Pfeiffer et al., 2008; Tirian and Dickson, 2017) to generate color depth MIPs. The Janelia and ~3000 of VT brain images and VNCs were imaged by the FlyLight project team (https://www.janelia.org/project-team/flylight). The rest of the VT brain images were imaged in Vienna (Tirian and Dickson, 2017). The Janelia collection brains were aligned by JBA and CMTK (Peng et al., 2011; Rohlfing and Maurer, 2003). VT brains are aligned by Amira software then transformed to JFRC2010 template space by CMTK. The color depth MIPs are created by our new program based on Fiji (https://fiji.sc/) https://www.janelia.org/open-science/color-depth-mip. All of the VNC (15,057 of Gen1-GAL4 and 3,534 of LexA) are imaged by the FlyLight project. All VNC are aligned by our newly coded VNC global aligner + CMTK.

## Acknowledgement

We thank Gerry Rubin, Barry Dickson and the other members of the FlyLight steering committee for supporting the Color MIP tool development and VNC aligner. Wyatt Korff and Robert Svirskas for the editing and discussion of the manuscript. We thank Robert Court for an early version and inspiration for the VNC aligner. The FlyLight Project Team for the generation of original confocal files. Sean Murphy for encouragement during this project.

